# Aboveground biomass increments over 26 years (1993–2019) in an old-growth cool-temperate forest in northern Japan

**DOI:** 10.1101/2021.07.02.450668

**Authors:** Mahoko Noguchi, Kazuhiko Hoshizaki, Michinari Matsushita, Daiki Sugiura, Tsutomu Yagihashi, Tomoyuki Saitoh, Tomohiro Itabashi, Ohta Kazuhide, Mitsue Shibata, Daisuke Hoshino, Takashi Masaki, Katsuhiro Osumi, Kazunori Takahashi, Wajirou Suzuki

**Affiliations:** Tohoku Research Center, Forestry and Forest Products Research Institute, Morioka, 020-0123, Japan; Department of Biological Environment, Akita Prefectural University, Akita, 010-0195, Japan; Forest Tree Breeding Center, Forestry and Forest Products Research Institute, Hitachi, 319-1301, Japan; Forestry and Forest Products Research Institute, Tsukuba, 305-8687, Japan; Field Science Center, Faculty of Agriculture, Tottori University (retired), Tottori, 680-8553, Japan; Kansai Research Center, Forestry and Forest Products Research Institute, Kyoto, 612-0855, Japan; Forestry and Forest Products Research Institute (retired), Tsukuba, 305-8687, Japan

**Keywords:** forest biomass, long-term data, Kanumazawa Riparian Research Forest, temperature

## Abstract

Assessing long-term changes in biomass of old-growth forests is critical in evaluating forest ecosystem functions under a changing climate. Long-term biomass changes are the result of accumulated short-term changes, which can be affected by endogenous processes such as gap filling in small-scale canopy openings. Here, we used 26 years (1993–2019) of repeated tree census data in an old-growth, cool-temperate, deciduous mixed forest that contains three topographic units (riparian, denuded slope, and terrace) in northern Japan to document decadal changes in aboveground biomass (AGB) and their processes in relation to endogenous processes and climatic factors. AGB increased steadily over the 26 years in all topographic units, but different tree species contributed to the increase among the topographic units. AGB gain within each topographic unit exceeded AGB loss via tree mortality in most of the measurement periods despite substantial temporal variation in AGB loss. At the local scale, variations in AGB gain were partially explained by compensating growth of trees around canopy gaps. Climate affected the local-scale AGB gain: the gain was larger in the measurement periods with higher mean temperature during the current summer but smaller in those with higher mean temperature during the previous autumn, synchronously in all topographic units. The decadal climate trends of warming are likely to have contributed to the steady increase in AGB in this old-growth forest.

## Introduction

Old-growth forests are widely recognized to play an important role in the carbon cycle (Luyssaert et al. 2008). It has been commonly accepted that old-growth forests are carbon neutral (Odum 1969) and their aboveground biomass (AGB) is at ‘steady state’, with equal gross primary production and respiration (Bormann and Likens 1979). However, recent studies indicate that they grow continuously (Foster et al. 2014) and work as carbon sinks with increasing biomass over centuries (Luyssaert et al. 2008). As biomass growth of old-growth forests is more susceptible to climate change than that in young forests (Chen et al. 2016), assessing long-term changes in biomass of old-growth forests is critical in evaluating the effects of climate change on forest ecosystem functions (McDowell et al. 2020).

Long-term changes in biomass result from the accumulation of short-term changes in the form of gain due to tree growth and loss due to mortality (Hoshizaki et al. 2004). Therefore, to understand how climate affects changes in AGB, the effects of climatic factors on each component need to be taken into account (Chen and Luo 2015; Peña et al. 2018). In addition, endogenous processes such as gap filling in small-scale canopy openings can drive biomass change (Phillips et al. 2009; McDowell et al. 2020): at the local scale, gap formation may cause first a decrease and then an increase in AGB caused by growth promotion of trees around the gap. Repeatedly measured tree census data with tree location can be useful in revealing these processes.

Environmental factors such as topographic position affect both forest biomass (Kubota et al. 2004; Valencia et al. 2009) and tree species composition (Chen and Luo 2015; Kuuluvainen et al. 2017; Ohmann and Spies 1998). For instance, on northern Honshu, Japan, *Fagus crenata* often dominates forest stands on hillslopes, whereas more tree species occur in riparian areas (Suzuki et al. 2002). Tree species in riparian forests have diverse life history traits (e.g., both shorter and longer lifespans, heavy sprouting; Nakamura and Inahara 2007; Sakio 2020). Therefore, hillslope and riparian stands are expected to differ in the dynamics (i.e., growth and mortality) and, consequently, the pattern of biomass changes in component species. In addition, a recent analysis of long-term tree census data in northern Japan has revealed different responses among species to changing climate and consequent changes in stand structure and species composition (Hiura et al. 2019). Thus, stands with different topographic characteristics can show different responses to climate change.

Here, we quantify decadal changes in AGB and their processes in relation to endogenous processes and climatic factors, using tree census data measured repeatedly over 26 years (1993–2019) in an old-growth, cool-temperate deciduous mixed forest with different types of topographic units in northern Japan. We ask the following questions: (1) Did AGB show net increase or decrease over the whole forest and study period? (2) Did tree species contribute differently to biomass change among the different types of topographic units? (3) How did gain and loss contribute to the overall changes in stand biomass? (4) Did climatic factors and endogenous processes such as canopy gap formation and recovery influence short-term changes in AGB at the local scale?

## Materials and methods

### Study site

The study was conducted in the Kanumazawa Riparian Research Forest (KRRF) in Iwate, northern Japan (39°06′N, 140°51′E), an old-growth forest with no record of significant anthropogenic disturbances. In KRRF, tree community dynamics have been repeatedly measured since the establishment of a 4.71-ha permanent plot in 1993 (Fig. 1; Suzuki et al. 2002). This site is one of the core research sites of the Japan Long-Term Ecological Research Network (JaLTER). KRRF has a cool-temperate climate with a mean annual temperature of 8.8 °C and a warmth index (Kira 1991) of 71 °C month (Noguchi et al. unpublished). The mean annual precipitation is 2000 mm, and snow cover lasts 5 months with maximum snow depth of approximately 2 m (Oki et al. 2013). The vegetation depends on the topographic unit. The riparian area is covered with a species-rich deciduous broadleaved forest consisting of both riparian specialists (*Cercidiphyllum japonicum*, *Aesculus turbinata*, *Acer mono*, *Pterocarya rhoifolia*, and *Ulmus laciniata*) and habitat generalists (*Fagus crenata* and *Quercus crispula*) (Masaki et al. 2008; Suzuki et al. 2002). The upper slopes and terrace are dominated by *F. crenata* and *Q. crispula*. Detailed information on the ecology of component species is available in Hoshizaki et al. (1997, 1999), Masaki et al. (2005), and Osumi (2006). The age of the largest *C. japonicum* individual is estimated to be more than 500 years (Osumi 2006), indicating that this forest is sufficiently old-growth. The natural disturbance regime in KRRF is characterized by canopy gap formation and fluvial sediment movements (Oki et al. 2013). Gap-creating disturbance occurs about every 1 to 3 years, with gap size ranging from tens to hundreds of square meters (Oki et al. 2013). Recent fluvial sediment movements were recorded in 1988, 1998, and 2007, causing ground disturbance with sizes ranging from 144 m^2^ to 680 m^2^ but no damage to canopy trees.

**Fig. 1.**
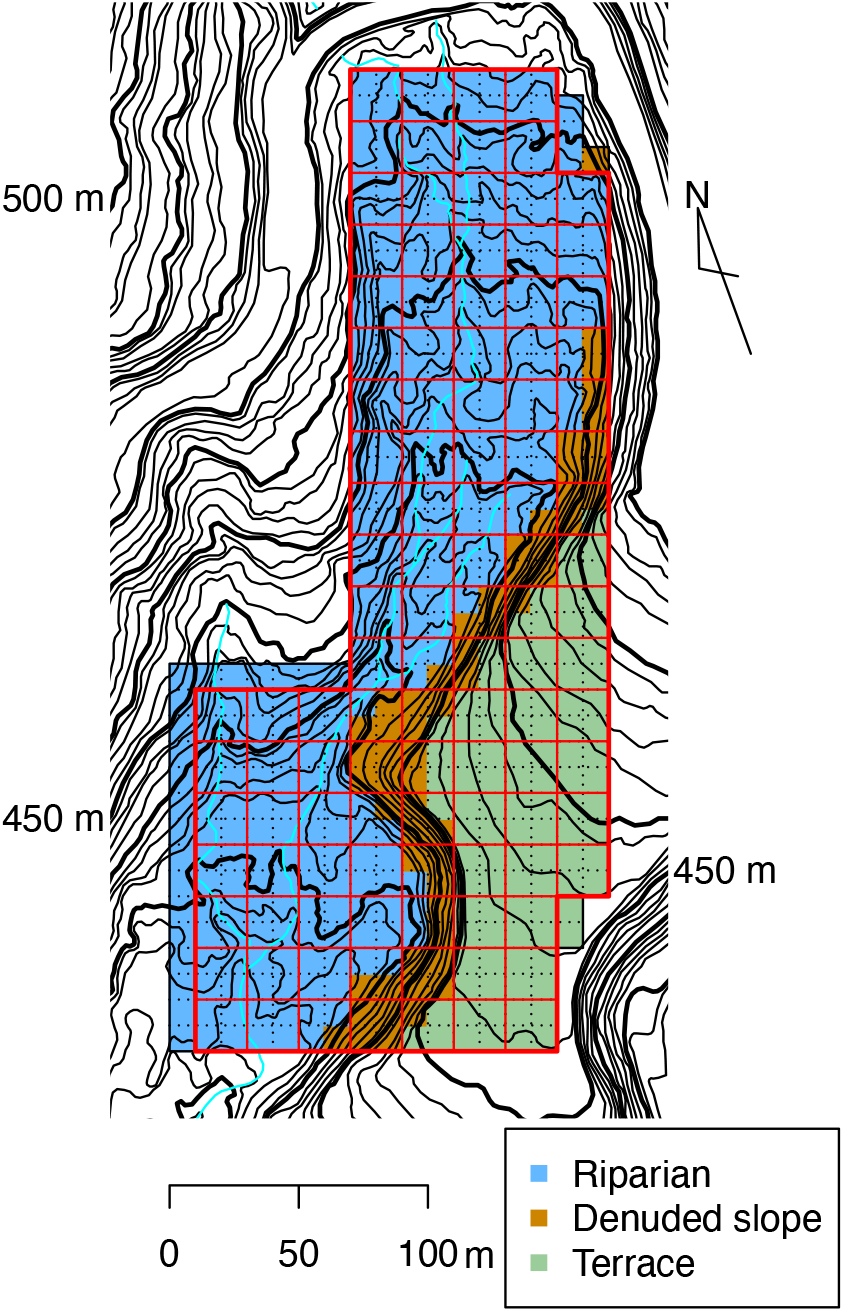
Topographic map of the Kanumazawa Riparian Research Forest (KRRF). The solid frame represents the 4.71-ha KRRF plot. Colors denote the three topographic units: blue, riparian (3.11 ha); orange, denuded slope (0.57 ha); green, terrace (1.06 ha). Black dotted lines show the 10-m × 10-m quadrats; thin red lines show the 20-m × 20-m subplots. Contour interval is 2 m.

### Field measurement

The 4.71-ha permanent plot was divided to 471 10-m × 10-m quadrats (Fig. 1). The plot ranges in elevation from 400 to 460 m a.s.l., and includes three topographic units: riparian (3.11 ha), terrace (1.06 ha), and denuded slope between them (0.57 ha). In the whole plot, all stems greater than 5 cm in diameter at breast height (DBH) were tagged for identification and mapped, and DBH was measured at the same marked location on each stem every 2 years from 1993 to 1999 and every 4 years from then to 2019.

### Estimation of AGB change and its components

We calculated tree AGB, basal area (BA), and stem density in each topographic unit. Individual tree AGB was estimated by using a general allometric equation for tree species in Japan (Ishihara et al. 2015):

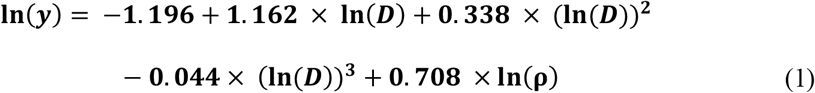

where *y* is AGB; *D* is stem DBH, and ρ is the wood specific gravity of each species (Editorial Board of Wood Industry 1966; Kurokawa et al. unpublished; Fujiwara et al. 2007). Confidence intervals of changes in AGB, BA, and stem density were estimated via bootstrapping across 10-m × 10-m quadrats following the method of Valencia et al. (2009).

To overview trends in AGB change during the study period and net annual change in AGB, we calculated AGB for three tree size classes: large (≥50 cm DBH), medium (15–50 cm DBH), and small (5–15 cm DBH). The net annual change in AGB (in Mg ha^−1^ y^−1^) was calculated each 4-year period from 1996 to 2019 from the tree DBH data of 1995 to 2019. It was then dissected into annual AGB gain and annual AGB loss. Annual AGB gain was calculated separately for growth of trees in each size class and ingrowth, and AGB loss was calculated for mortality of trees in each size class.

### Analysis of factors affecting short-term AGB gain at local scale

We examined the effects of climatic condition in each measurement period, canopy gap formation and topography on local-scale AGB gain using two generalized linear mixed-effect models (GLMMs). For both models, variables were calculated for every 20-m × 20-m subplot in each 4-year period from 1996 to 2019. Subplot size was determined as an appropriate area to detect gap formation and subsequent recovery in consideration of the range of gap size in KRRF. Both models used AGB gain as the response variable and included subplot as a random effect.

In model 1, we aimed to investigate whether the amount of AGB gain differed among the measurement periods with the effects of topographic unit and gap formation in the current and previous measurement periods. Fixed effects were initial AGB in the current measurement period, AGB losses in the current and previous measurement periods, topographic unit, and the five 4-year measurement periods between 2000 and 2019, with topographic unit and measurement period as categorical variables. Initial AGB was included as it is expected to be the “capital” for AGB gain by tree growth. AGB losses were indices of gap formation in the current and previous measurement periods.

In model 2, the effect of climate was analyzed separately from the effect of measurement period to avoid multicollinearity. Fixed effects were initial AGB in the current measurement period, AGB loss in the current and previous measurement periods, topographic unit, and mean temperature during the previous autumn (September–November) and the current summer (June–August) over the measurement periods. Both types of mean temperature have a major influence on annual DBH growth of individual trees in most dominant species of KRRF (Matsushita et al. manuscript in preparation). As the on-site temperature data do not cover the entire study period, we used data from the nearest weather station, at Wakayanagi (39°08′N, 141°04′E; 97 m a.s.l.: Japan Meteorological Agency, https://www.data.jma.go.jp/gmd/risk/obsdl/index.php), 18 km east of the study site. These models were fitted by the lme4 v. 1.1-21 package (Bates et al. 2015) in R 3.6.3 (R Core Team 2020). To evaluate the variance explained by the models, we calculated two *R*^2^ values for mixed-effect models following the method of Nakagawa et al. (2017): marginal *R^2^* (*R*^2^_GLMM(m)_), which is the proportion of the variance explained by fixed effects, and conditional *R^2^* (*R*^2^_GLMM(c)_), which is the proportion of the variance explained by both fixed and random effects. These were calculated by the MuMIn v. 1.43.15 package (Bartoń 2019) in R.

## Results

### Overall changes in AGB at plot scale

BA in 1993 was greatest in the riparian unit (34.2 m^2^ ha^−1^) and least in the denuded slope unit (Table 1). From 1993 to 2019, BA increased significantly in all topographic units. AGB was greatest in the terrace unit (246.0 Mg ha^−1^) at the beginning of the study period (Table 1). It increased significantly in all topographic units during the study period, increasing in most 4-year periods except for some short pauses; for instance, from 2011 to 2015 in the riparian and denuded slope units (Fig. 2). AGB of large trees (≥50 cm DBH) in 1993 occupied 76.7% of total AGB in riparian, 70.6% in denuded slope, and 77.7% in terrace units. Trends of increasing total AGB in the riparian and terrace units were similar to those of large-tree AGB. During the study period, stem density declined in the riparian and terrace units but increased in the denuded slope unit (Table 1). The change in stem density was significant only in the riparian unit.

**Table 1.** Basal area, aboveground biomass, and stem density at the study site at the beginning (1993) and end (2019) of the study period with overall changes in three topographic units (riparian, denuded slope, and terrace).

|  | 1993 | 2019 | Change |
| --- | --- | --- | --- |
| Basal area (m<sup>2</sup> ha<sup>-1</sup>) |  |  |  |
| Riparian | 34.2 (28.7–40.5) | 38.6 (32.4–457.7) | 4.5 (2.4–6.5) |
| Denuded slope | 21.5 (15.3–28.3) | 26.6 (20.2–33.8) | 5.1 (2.2–7.8) |
| Terrace | 32.3 (26.6–37.7) | 32.3 (26.6–37.7) | 4.3 (1.5–6.7) |
| Aboveground biomass (Mg ha<sup>-1</sup>) |  |  |  |
| Riparian | 244.1 (202.7–289.4) | 274.2 (230.1–326.2) | 30.1 (14.0–45.6) |
| Denuded slope | 156.8 (100.3–216.0) | 191.5 (136.1–252.7) | 34.7 (13.2–56.1) |
| Terrace | 246.0 (202.4–293.5) | 276.7 (225.8–336.4) | 30.6 (8.0–53.4) |
| Stem density (stems ha<sup>-1</sup>) |  |  |  |
| Riparian | 583 (519–648) | 509 (452–581) | –73 (–111 to –36) |
| Denuded slope | 781 (637–939) | 877 (704–1046) | 96 (–35 to 235) |
| Terrace | 952 (833–1061) | 906 (785–1031) | –47 (–118 to 24) |

Values in parentheses are 95% confidence intervals. When CIs do not include 0, the changes are significant.

**Fig. 2.**
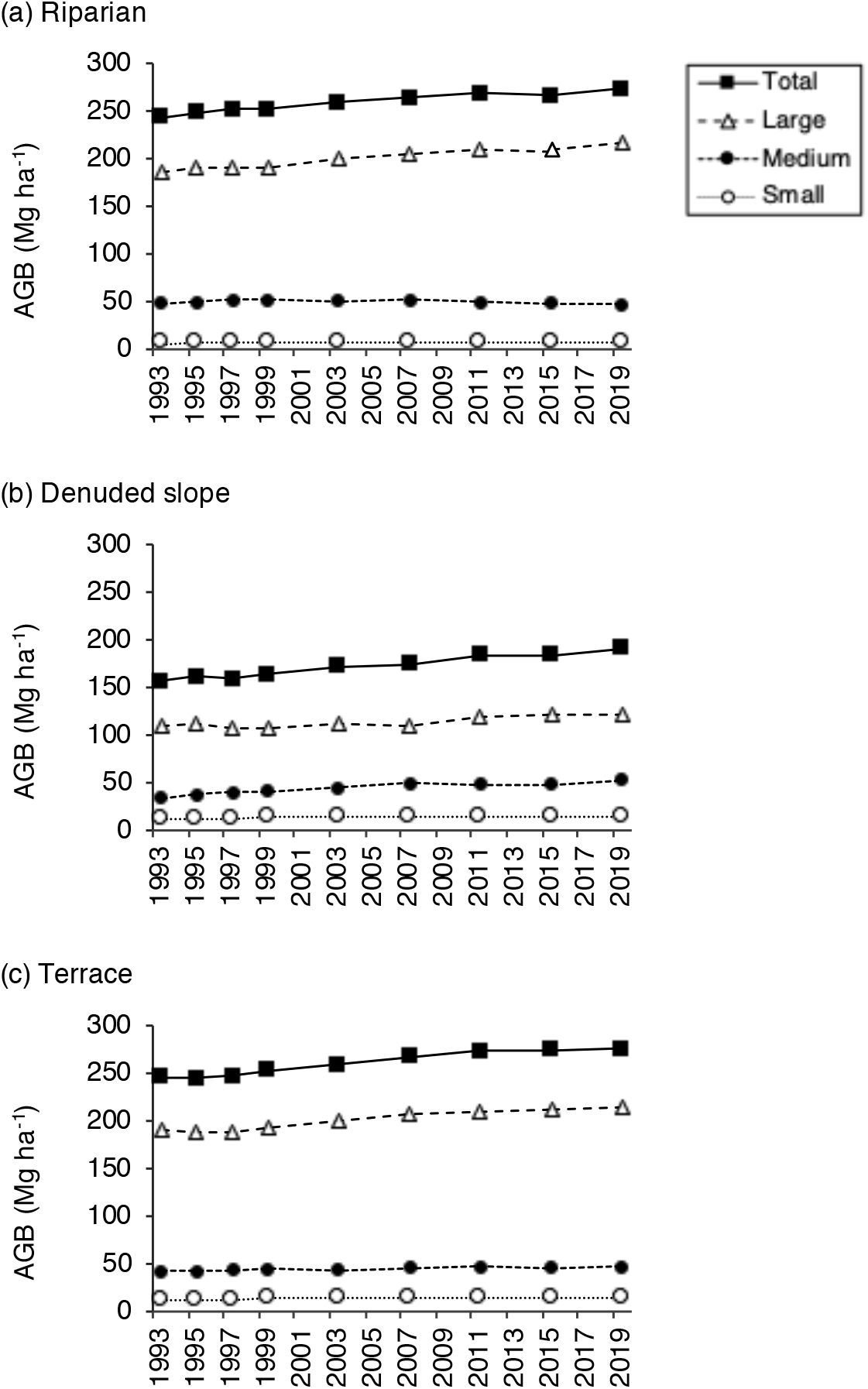
Trends in total aboveground biomass (AGB) over 26 years in the three topographic units. AGB is shown for stand total and stems in three size classes: large (diameter at breast height [DBH] ≥ 50 cm), medium (DBH, 15–50 cm), and small (DBH, 5–15 cm).

In the riparian unit, *C. japonicum* had the largest AGB at the beginning of the study period, followed by *F. crenata*, *A. turbinata*, *Q. crispula*, and *A. mono* (Table 2). AGB of these species, except for *Q. crispula*, increased during the study period. *Pterocarya rhoifolia* had the greatest increment in AGB over the study period, accounting for 52.3% of the total increment in the riparian unit, followed by *A. turbinata* at 25.4%. In contrast, several other species with relatively small AGB at the beginning, such as *Zelkova serrata* and *Ulmus laciniata*, showed a decline in AGB during the study period. The denuded slope and terrace units were dominated by *F. crenata* and *Q. crispula*, and the denuded slope by *A. mono* as well (Table 2). All these species had an increase in AGB during the study period, maintaining the AGB-based rank of species composition.

**Table 2.** Overall changes in aboveground biomass (AGB, in Mg ha^−1^) of component tree species in each topographic unit during the study period and the relative contribution of each species to the total change in AGB.

| Species | Riparian |  |  |  | Denuded slope |  |  |  | Terrace |  |  |  |
| --- | --- | --- | --- | --- | --- | --- | --- | --- | --- | --- | --- | --- |
|  | 1993 | 2019 | Change | Contribution | 1993 | 2019 | Change | Contribution | 1993 | 2019 | Change | Contribution |
|  |  |  |  | (%) |  |  |  | (%) |  |  |  | (%) |
| *Fagus crenata* | 52.1 | 56.2 | 4.1 | 13.7 | 68.9 | 80.4 | 11.5 | 33.2 | 139.3 | 153.9 | 14.7 | 47.9 |
| *Quercus crispula* | 30.6 | 30.3 | -0.2 | -0.8 | 30.4 | 37.8 | 7.4 | 21.2 | 82.1 | 86.3 | 4.2 | 13.7 |
| *Cercidiphyllum japonicum* | 58.9 | 61.9 | 3.0 | 10.1 | 0.0 | 0.0 | 0.0 | 0.0 | 0.0 | 0.0 | 0.0 | 0.0 |
| *Aesculus turbinata* | 47.8 | 55.5 | 7.6 | 25.4 | 3.8 | 5.4 | 1.6 | 4.5 | 0.0 | 0.0 | 0.0 | 0.0 |
| *Acer mono* | 21.0 | 21.6 | 0.5 | 1.8 | 28.9 | 38.0 | 9.0 | 26.1 | 0.2 | 0.4 | 0.2 | 0.6 |
| *Pterocarya rhoifolia* | 8.6 | 24.4 | 15.8 | 52.4 | 0.9 | 3.7 | 2.8 | 8.0 | 0.0 | 0.0 | 0.0 | 0.0 |
| *Zelkova serrata* | 5.7 | 4.7 | -1.0 | -3.2 | 2.0 | 3.6 | 1.5 | 4.4 | 0.0 | 0.0 | 0.0 | 0.0 |
| *Ulmus laciniata* | 4.8 | 2.8 | -2.0 | -6.6 | 0.0 | 0.0 | 0.0 | 0.0 | 0.0 | 0.0 | 0.0 | 0.0 |
| *Magnolia obovata* | 4.2 | 6.0 | 1.8 | 6.1 | 0.0 | 0.5 | 0.4 | 1.3 | 1.2 | 2.2 | 1.0 | 3.3 |
| *Kalopanax pictus* | 4.5 | 5.0 | 0.5 | 1.8 | 0.1 | 0.7 | 0.6 | 1.7 | 0.0 | 0.3 | 0.3 | 0.9 |
| *Acer sieboldianum* | 0.4 | 0.1 | -0.3 | -1.1 | 4.5 | 3.5 | -1.0 | -2.9 | 6.0 | 8.0 | 2.0 | 6.5 |
| *Acer japonicum* | 0.5 | 0.9 | 0.4 | 1.3 | 1.9 | 2.4 | 0.5 | 1.5 | 6.0 | 7.4 | 1.4 | 4.5 |
| Others | 5.0 | 4.8 | -0.2 | -0.8 | 15.3 | 15.7 | 0.4 | 1.1 | 11.2 | 18.1 | 6.9 | 22.5 |
| Total | 244.1 | 274.2 | 30.1 | 100.0 | 156.8 | 191.5 | 34.7 | 100.0 | 246.0 | 276.7 | 30.6 | 100.0 |

Annual gain in AGB remained at approximately 3 Mg ha^−1^ y^−1^ with some differences among the measurement periods: larger in 2008–2011 and 2016–2019 and smaller in 2004– 2007 in all topographic units (Fig. 3). In the riparian and terrace units, large- and medium-sized trees accounted for most of the annual gain. Annual losses in AGB fluctuated among the 4-year periods, and were largest in 2012–2015 in all topographic units. Regardless of topographic unit, measurement periods with greater loss of AGB of large trees tended to have greater total loss of AGB. As a consequence, net annual change in AGB ranged from −0.6 to +2.6 Mg ha^−1^ y^−1^ but stayed positive except in 2012 to 2015 in the riparian and denuded slope units.

**Fig. 3.**
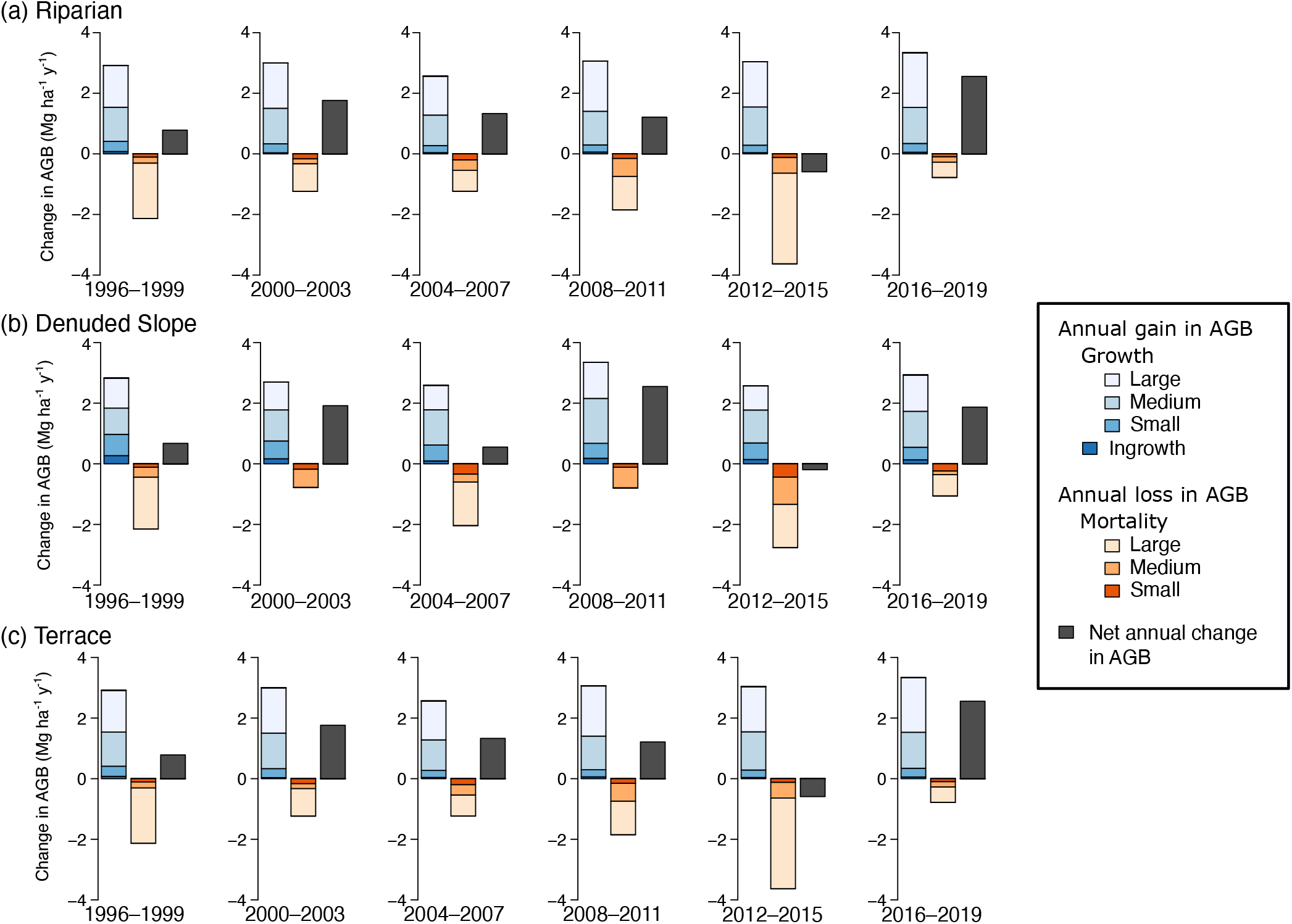
Components of average annual change in aboveground biomass (AGB) by each measurement period in the three topographic units. Blue bars (4 levels of color gradient) denote annual AGB gain from growth of surviving stems in the three size classes (large, diameter at breast height [DBH] ≥ 50 cm; medium, 15– 50 cm DBH; small, 5–15 cm DBH) and ingrowth. Orange bars (3 levels of color gradient) denote annual AGB loss from stems that died during each measurement period in the three size classes. Dark gray bars denote net average annual change in AGB.

### Effects of climate and gap formation on short-term AGB gain at local scale

Local-scale AGB gain in the 20-m × 20-m subplots was positively influenced by initial AGB in each measurement period (GLMM model 1: Table 3, Fig. 4). It was significantly greater in subplots with larger AGB loss in the previous measurement period but smaller in subplots with larger AGB loss in the current measurement period. It was not significantly affected by topographic unit. It differed significantly among the measurement periods: smaller in 2004– 2007 and larger in 2008–2011 and 2016–2019. In model 1, *R*^2^_GLMM(m)_ = 0.31 and *R*^2^_GLMM(c)_ = 0.76, indicating that 31% of the variation was explained by fixed effects and 76% by fixed and random effects. In model 2 (Table 4), the effects of initial AGB, AGB loss in the current and previous measurement periods, and topographic unit were almost identical to those in model 1. Local-scale AGB gain was larger in measurement periods with higher mean temperature during the current summer but smaller in those with higher mean temperature during the previous autumn. Model 2 explained almost identical variation as model 1, with *R*^2^_GLMM(m)_ = 0.32 and *R*^2^_GLMM(c)_ = 0.77.

**Fig. 4.**
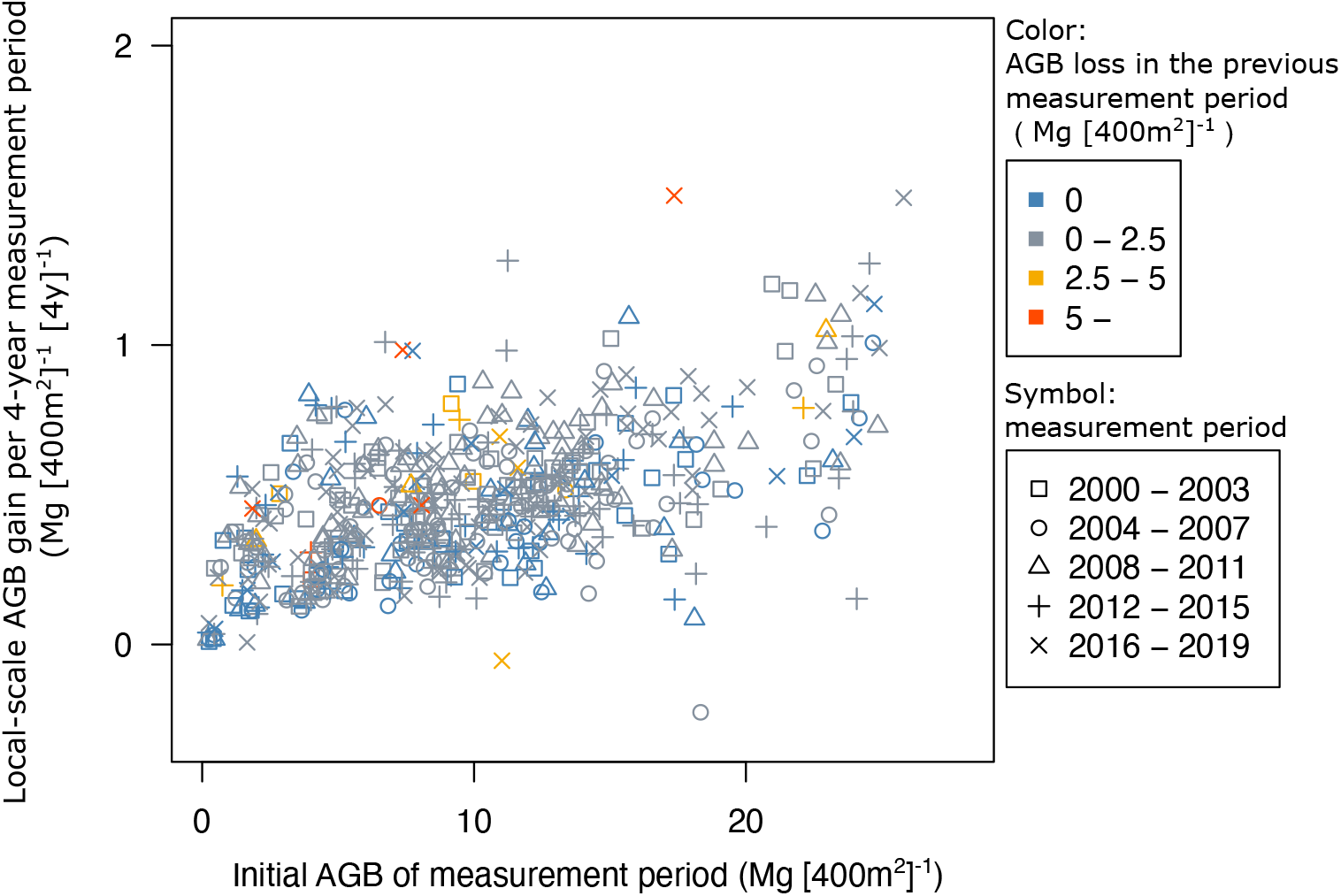
Local-scale aboveground biomass (AGB) gain per 4-year measurement period in relation to initial AGB of measurement period. Colors represent classes of AGB loss in previous measurement period; symbols represent measurement periods.

**Table 3.**
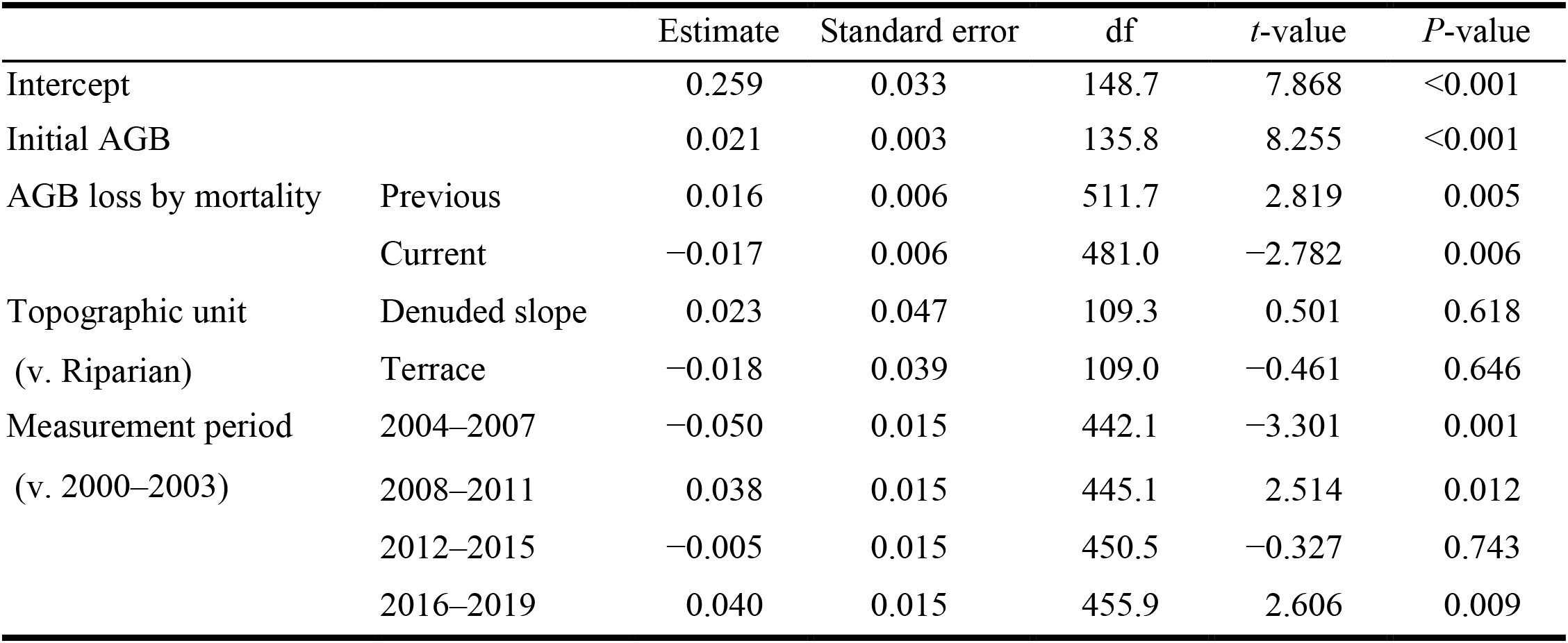
Results of the generalized linear mixed-effect model (model 1) testing the effects of initial aboveground biomass (AGB), canopy gap formation, topographic unit, and measurement period on the AGB gain in 20-m × 20-m subplots.

|  |  | Estimate | Standard error | df | *t*-value | *P*-value |
| --- | --- | --- | --- | --- | --- | --- |
| Intercept |  | 0.259 | 0.033 | 148.7 | 7.868 | <0.001 |
| Initial AGB |  | 0.021 | 0.003 | 135.8 | 8.255 | <0.001 |
| AGB loss by mortality | Previous | 0.016 | 0.006 | 511.7 | 2.819 | 0.005 |
|  | Current | −0.017 | 0.006 | 481.0 | −2.782 | 0.006 |
| Topographic unit<br>(v. Riparian) | Denuded slope | 0.023 | 0.047 | 109.3 | 0.501 | 0.618 |
|  | Terrace | −0.018 | 0.039 | 109.0 | −0.461 | 0.646 |
| Measurement period<br>(v. 2000–2003) | 2004–2007 | −0.050 | 0.015 | 442.1 | −3.301 | 0.001 |
|  | 2008–2011 | 0.038 | 0.015 | 445.1 | 2.514 | 0.012 |
|  | 2012–2015 | −0.005 | 0.015 | 450.5 | −0.327 | 0.743 |
|  | 2016–2019 | 0.040 | 0.015 | 455.9 | 2.606 | 0.009 |

**Table 4.** Results of the generalized linear mixed-effect model (model 2) testing the effects of initial aboveground biomass (AGB), canopy gap formation, topographic unit, and climate (mean air temperature) of each measurement period on the AGB gain in 20-m × 20-m subplots.

|  |  | Estimate | Standard error | df | *t*-value | *P*-value |
| --- | --- | --- | --- | --- | --- | --- |
| Intercept |  | −0.559 | 0.321 | 445.1 | −1.738 | 0.083 |
| Initial AGB |  | 0.021 | 0.003 | 137.9 | 8.294 | <0.001 |
| AGB loss by mortality | Previous | 0.016 | 0.006 | 508.0 | 2.925 | 0.004 |
|  | Current | −0.015 | 0.006 | 481.7 | −2.550 | 0.011 |
| Topographic unit<br>(v. Riparian) | Denuded slope | 0.023 | 0.047 | 109.4 | 0.498 | 0.619 |
|  | Terrace | −0.018 | 0.038 | 109.1 | −0.462 | 0.645 |
| Mean air temperature | Previous autumn | −0.137 | 0.023 | 443.0 | −5.865 | <0.001 |
|  | Current summer | 0.120 | 0.021 | 450.8 | 5.803 | <0.001 |

## Discussion

AGB of KRRF increased steadily over the 26 years in all topographic units, with increments of 30 to 35 Mg ha^−1^ in each (Table 1). BA also increased over the study period, even though it was initially equivalent to values reported in other cool-temperate old-growth forests in Japan (Masaki et al. 1992; Nakashizuka 1988; Seiwa et al. 2013), indicating that the forest had already been well stocked. These results are consistent with reports that temperate old-growth forests continuously gain biomass over the long term (Keeton et al. 2011; Luyssaert et al. 2008). This continuous stand-scale biomass increment was attributable mainly to an increase in AGB of large trees, in agreement with the reported global importance of large trees in determining stand AGB (Lutz et al. 2018; Slik et al. 2013).

Patterns of tree growth or stand-biomass-change vary across tree species composition and diversity, as well as with environmental conditions such as topography (Kubota et al. 2004; Valencia et al. 2009). In KRRF, topography was reported to determine tree species distribution through affecting seedling survival differently across species (Masaki et al. 2005). Here, however, a steady increase in AGB was common to all three topographic units (Table 1; Fig. 2), despite the difference in tree species composition among them (Table 2). In addition, local-scale AGB gain did not differ among the topographic units (Table 3). We attribute this similarity to the distinct AGB increment in *F. crenata*, which is dominant in all three topographic units (Table 2). A growing abundance of *F. crenata* is documented in several stable old-growth forest (Seiwa et al. 2013; Yamamoto and Nishimura 1999). Increases in both AGB and stem density of *F. crenata* in KRRF may be due to lack of remarkable disturbance even in the riparian unit during the study period. *Pterocarya rhoifolia*, a riparian specialist of cool-temperate forests in Japan (Sakio et al. 2002), made the largest contribution to the stand AGB increment in the riparian unit (Table 2). Despite the substantial decline in its stem density (Appendix 1), its AGB at the end of the study period was 3 times the initial value. It is likely that the fast growth of *P. rhoifolia* (Sakio 1993) is associated with its rapid increase in AGB. Although AGB decreased in some species such as *Z. serrata* and *U. laciniata* in the riparian unit, *P. rhoifolia* compensated for the decrease and resulted in the stand-level AGB increase.

Mortality is the major cause of reduced growth or decline in AGB (Schuster et al. 2008; Xu et al. 2012). Although large disturbances such as strong typhoons, insect outbreaks, or severe flooding in the riparian unit were not recorded during the 26 years, the loss of AGB varied substantially among the topographic units and among the 4-year measurement periods (Fig. 3). These variations were explained mainly by the spatio-temporal variation in mortality of large trees. A significant contribution of large-tree mortality to the AGB loss has also been reported in other old-growth forests (Hoshizaki et al. 2004). Despite these temporal and spatial variations, the AGB loss generally remained smaller than the AGB gain, bringing about a positive change in AGB in most of the measurement periods. The temporal change of stand-level AGB appears to be inconsistent with the assumed long-term balance between biomass loss caused by canopy gap formation and subsequent gain during gap recovery.

In contrast to AGB loss, temporal fluctuations in AGB gain were generally synchronized across the topographic units at the stand scale (Fig. 3). The results of model 1 indicate that local-scale AGB gain also differed among measurement periods even after adjustment for initial AGB and disturbance during each period (Table 3). As expected, initial AGB positively influenced local-scale AGB gain (Fig.4). Larger AGB loss in the previous measurement period caused greater AGB gain, suggesting that variations in local-scale AGB gain are partially explained by recovery in and around canopy gaps. Local-scale AGB gain also substantially differed among the measurement periods. The results of model 2 suggest that the observed temporal variations in AGB gain are caused by climatic factors: a hotter current summer had positive effects whereas a warmer previous autumn had negative effects on AGB gain (Table 4). The positive response to high temperature in the growing season is consistent with the trends of individual-tree growth observed in deciduous broadleaved species in KRRF (Matsushita et al. manuscript in preparation) and other cool-temperate forests in Japan (Hiura et al. 2019). The negative effect of a warmer autumn on growth is also found in individual-tree growth in KRRF (Matsushita et al. manuscript in preparation) and may be caused by a larger increment in respiration than in photosynthesis (Piao et al. 2008).

Our models explained a considerable amount of variation in local AGB gain, although the analysis did not include other potential factors that enhance tree growth such as change in precipitation (Hiura et al. 2019), rising CO_2_ levels in the atmosphere (e.g., Norby et al. 2005), and nitrogen deposition (Thomas et al. 2010). The temperature at the weather station nearest to KRRF shows a substantial rise in summer and autumn temperatures over the past 40 years (Appendix 2). Decadal climate trends of warming are likely to have contributed to the observed steady increase in AGB in KRRF by providing favorable conditions for tree growth in the study period. Warming-induced growth acceleration in the past several decades has been reported in temperate and boreal forests in Europe (Kauppi et al. 2014; Pretzsch et al. 2014). In Japan, some cool-temperate beech-dominated forests are predicted to be vulnerable to warming (Matsui et al. 2009). Accumulation of tree growth data from a broader range of temperatures across cool-temperate forests in Japan will improve our understanding of the influence of climate change on this type of forest.

## Acknowledgements

We thank many colleagues, including Gaku Hitsuma, Chinatsu Homma, Shoji Naoe, Takayuki Ota and Takanobu Yagi for field assistance. We also thank Haruko Narita and Saori Sato for data preparation. Part of the data of this study was obtained from the Monitoring Sites 1000 Project of the Ministry of the Environment, Japan. This study was partly supported by JSPS KAKENHI Grant Numbers 15H04517, 19H02999 and 21H04946. We have no conflict of interest to declare.

