## Appendix 1 and 2 for "Aboveground biomass increments over 26 years (1993–2019) in an old-growth cool-temperate forest in northern Japan"

Appendix 1. Overall changes in stem density ($ha^{-1}$) of component tree species in each topographic unit during the study period. Species are listed in the same order as in Table 2 (i.e., the order of aboveground biomass [AGB] in 1993 in the entire plot).

| Species | Riparian unit | | | Denuded slope unit | | | Terrace unit | | |
|---|---|---|---|---|---|---|---|---|---|
| | 1993 | 2019 | Change | 1993 | 2019 | Change | 1993 | 2019 | Change |
| *Fagus crenata* | 68 | 90 | 22 | 107 | 146 | 39 | 87 | 91 | 4 |
| *Quercus crispula* | 15 | 11 | −4 | 44 | 46 | 2 | 43 | 41 | −2 |
| *Cercidiphyllum japonicum* | 86 | 82 | −4 | 0 | 0 | 0 | 0 | 0 | 0 |
| *Aesculus turbinata* | 42 | 39 | −3 | 4 | 5 | 2 | 0 | 0 | 0 |
| *Acer mono* | 36 | 39 | 3 | 68 | 60 | −9 | 1 | 1 | 0 |
| *Pterocarya rhoifolia* | 101 | 62 | −39 | 46 | 23 | −23 | 0 | 0 | 0 |
| *Zelkova serrata* | 13 | 10 | −2 | 5 | 5 | 0 | 0 | 0 | 0 |
| *Ulmus laciniata* | 40 | 24 | −16 | 2 | 2 | 0 | 0 | 0 | 0 |
| *Magnolia obovata* | 13 | 13 | 0 | 2 | 4 | 2 | 6 | 6 | 0 |
| *Kalopanax pictus* | 6 | 1 | −5 | 4 | 4 | 0 | 1 | 1 | 0 |
| *Acer sieboldianum* | 2 | 3 | 1 | 30 | 40 | 11 | 30 | 39 | 9 |
| *Acer japonicum* | 23 | 31 | 8 | 46 | 75 | 30 | 181 | 200 | 19 |
| Others | 136 | 103 | −33 | 425 | 468 | 44 | 604 | 527 | −77 |
| Total | 583 | 509 | −73 | 781 | 877 | 96 | 952 | 906 | −47 |

Appendix 2.

(a) Mean air temperature of the current summer

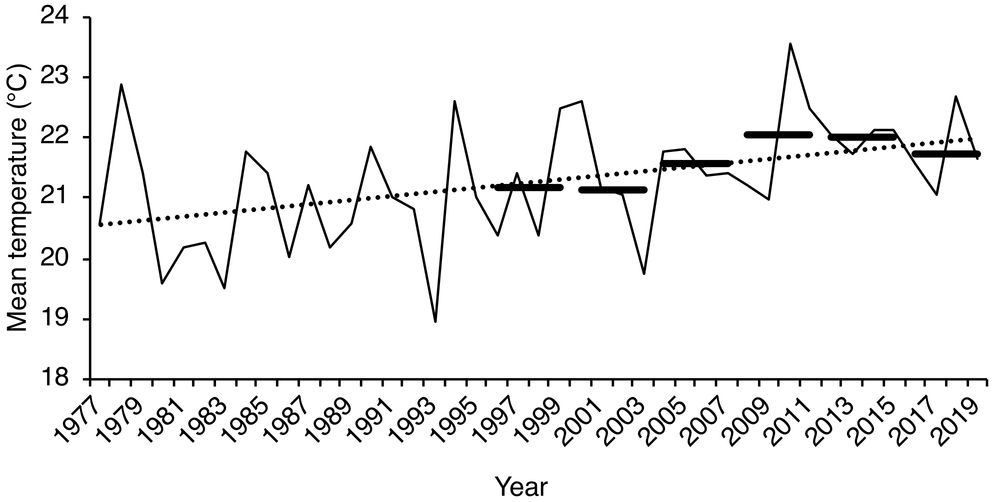


(b) Mean air temperature of the previous autumn

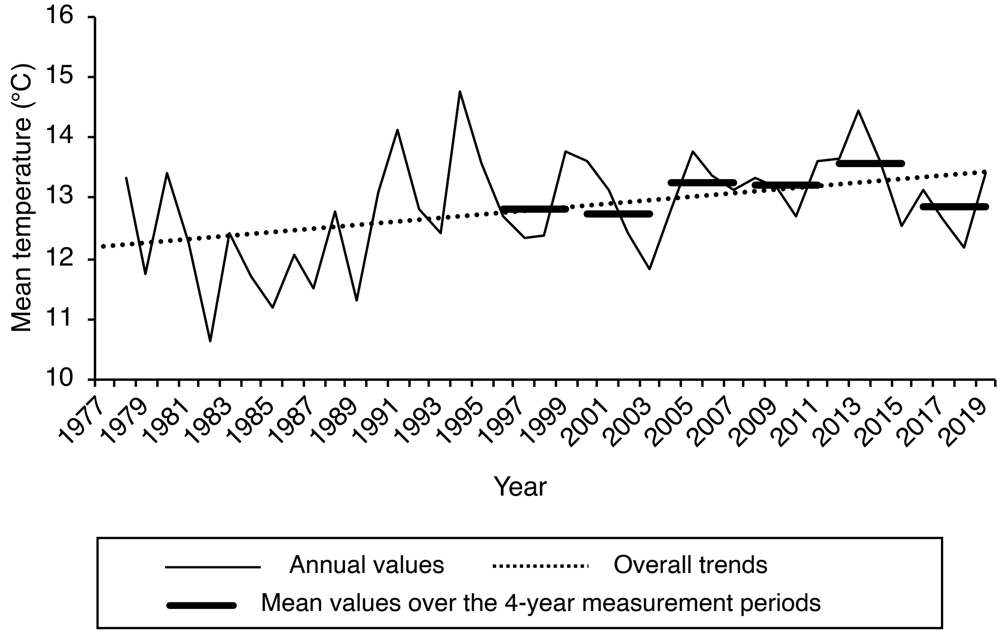


Trends in mean air temperature of (a) the current summer and (b) the previous autumn at the nearest weather station, Wakayanagi. Solid lines denote annual values; dotted lines denote overall trends. Solid horizontal bars denote the mean values over the six 4-year measurement periods.
